# Escalation of intravenous fentanyl self-administration and assessment of withdrawal behavior in male and female mice

**DOI:** 10.1101/2024.04.03.587958

**Authors:** Yueyi Chen, Tiange Xiao, Adam Kimbrough

**Author notes:** Correspondence: Dr. Adam Kimbrough, Department of Basic Medical Sciences, College of Veterinary Medicine, Purdue University, 625 Harrison Street, West Lafayette, IN, 47907.

## Abstract

**Background:** The rise in overdose deaths from synthetic opioids, especially fentanyl, necessitates the development of preclinical models to study fentanyl use disorder (FUD). While there has been progress with rodent models, additional translationally relevant models are needed to examine excessive fentanyl intake and withdrawal symptoms.

**Methods:** The study performed intravenous self-administration (IVSA) of fentanyl in male and female C57BL/6J mice for 14 days. Mechanical pain sensitivity during withdrawal was assessed using the von Frey test. Anxiety-like behavior was evaluated via the open field test one-week into abstinence and incubation of craving for fentanyl was assessed four weeks into abstinence.

**Results:** Both male and female mice demonstrated a significant escalation in fentanyl intake over the 14 days of self-administration, with significant front-loading observed in the final days of self-administration. Increased mechanical pain sensitivity was present from 36- to 48-hour into withdrawal and increased anxiety-like behavior was found 1 week into abstinence. Four weeks into abstinence, mice showed significantly higher active lever pressing than the final self-administration session prior to abstinence.

**Conclusions:** The study establishes a translationally relevant mouse model of IVSA of fentanyl, effectively encapsulating critical aspects of FUD, including escalation of drug intake, front-loading behavior, withdrawal symptoms, and prolonged craving for drug into abstinence. This model offers a robust basis for further exploration into behavioral and neurobiological mechanisms involved in fentanyl dependence and potential therapeutic strategies.

## Background

The opioid crisis has escalated into a severe public health issue, leading to over a hundred thousand overdose deaths annually, a trend sharply rising in recent years (1). Among various opioids, including heroin and prescription pain relievers like oxycodone and morphine, fentanyl stands out for its particularly devastating impact in recent years. Fentanyl, a synthetic opioid far more potent than its counterparts, has been implicated in the majority of these overdose deaths. In 2022, opioids were involved in 81.8% of overdose fatalities, with illegally made fentanyl accounting for 74.6% of these cases (2). The fentanyl crisis is not just about the sheer number of fatalities; it encompasses the complex behavioral and neural mechanisms of opioid dependence that need to be better understood.

Opioid use disorder (OUD) represents a relapsing disorder marked by an overwhelming desire to consume opioids, an inability to limit consumption, and the development of a significantly negative state both physically and emotionally when the drug is withdrawn (3). The *DSM-5* characterizes opioid use disorder as a problematic pattern of opioid use leading to clinically significant impairment or distress, as manifested by specific criteria such as a strong desire to use opioids, unsuccessful efforts to control use, and withdrawal symptoms when not using (4). Animal models are crucial for elucidating the etiology of OUD as they allow for the dissection of specific components of the cycle of OUD, including the development and intensification of drug-taking behavior and experience of withdrawal. An especially important component of animal models is high translational validity to human OUD. Rodent models using self-administration methods have revealed that a range of opioids can act as reinforcers, establishing stable self-administration behaviors. When given access to intravenous opioids, rodents develop a variety of somatic and motivational indicators of heroin dependence (5–7), oxycodone dependence (8–12), and morphine dependence (13, 14), that mirror the patterns of dependence seen in humans suffering from OUD. In those rodent models, there is a clear pattern of increasing drug intake over time, accompanied by intensified drug-seeking behavior. Withdrawal symptoms are also a hallmark of drug dependence and upon withdrawal from opioids, both rats and mice exhibit allodynia (15–18), along with anxiety-like behaviors (19), and a spectrum of other withdrawal-related physical symptoms (20–23). These observations highlight the translational validity and value of such models in deepening our comprehension of the behavioral and physiological changes associated with OUD.

However, many of the previous studies have focused on opioids other than fentanyl. Despite the growing need, current preclinical models of fentanyl dependence (e.g., fentanyl use disorder [FUD]) are limited, particularly in terms of simulating escalation of voluntary drug intake, especially over the course of multiple periods of drug use. Existing studies have predominantly focused on rat models, as evidenced by a growing body of literature (24–30). This is consistent across drugs (25, 31, 32), as common methods such as intravenous self-administration (IVSA) are substantially easier to perform in rats compared to mice. However, IVSA procedures have been used for mice successfully in a variety of drugs (6, 33–35) and mice offer unique advantages for studying the neurobiological mechanisms underlying FUD due to their viral and transgenic toolkits. Thus, it is critical to establish models of FUD in mice, including IVSA models. A previous study showed that mice are capable of initiating IVSA of fentanyl (33), and other mouse models have demonstrated withdrawal symptoms following oral or vapor self-administration (36–38).

A notable gap in the current scientific landscape is the absence of mouse models that effectively replicate the escalation of voluntary intravenous fentanyl consumption over prolonged periods and evaluate subsequent affective withdrawal phenomena. To address this gap, the present study sought to establish a robust and translationally valid IVSA of fentanyl model that results in significant withdrawal symptoms that can be used to study FUD. In the present study, we characterized the escalation of voluntary IVSA of fentanyl and subsequent signs of mechanical pain sensitivity and anxiety-like behavior, as well as incubation of fentanyl craving during abstinence in both male and female mice.

## Methods

### Experimental subjects

All of the experimental procedures were conducted in accordance with the guidelines of the National Institutes of Health (NIH) *Guide for the Care and Use of Laboratory Animals* and approved by the Purdue University Animal Care and Use Committee. Male and female C57BL/6J mice bred at Purdue University were used. Mice were between 9 to 11 weeks of age at the onset of the study. The animals were group housed, maintained on a 12/12 h light/dark cycle, and provided ad libitum access to food and water. Experimental procedures were carried out during the dark phase of the light/dark cycle.

### Drugs

Fentanyl citrate salt (Cayman Chemical, Ann Arbor, MI, USA, 22659) was dissolved in saline. Fentanyl solutions were filtered through 0.22 μm syringe filter for sterilization prior to use.

### Experiment Timeline

The experimental timeline is depicted in Figure 1. Mice underwent surgical implantation of jugular catheters into the right vein. Following a recovery period of 5-7 days, the mice were tested for baseline pain sensitivity and then underwent a 14-day period of daily sessions of IVSA of fentanyl. Following the 14 days, a progressive ratio test (PR) was performed, requiring the mice to press the active lever an exponentially increasing number of times to receive the drug (data provided in the supplementary materials). To allow for recovery to baseline drug intake levels, after the PR an additional day of standard IVSA session was performed. Mechanical pain sensitivity was assessed using the von Frey test at 12, 24, 36, and 48 hours into fentanyl withdrawal following the final IVSA session. An open field test was conducted one-week into fentanyl abstinence to evaluate changes in activity and anxiety-like behavior. Finally, four weeks into abstinence, mice were examined for active lever responses without drug infusion in an incubation of craving test.

**Figure 1.**
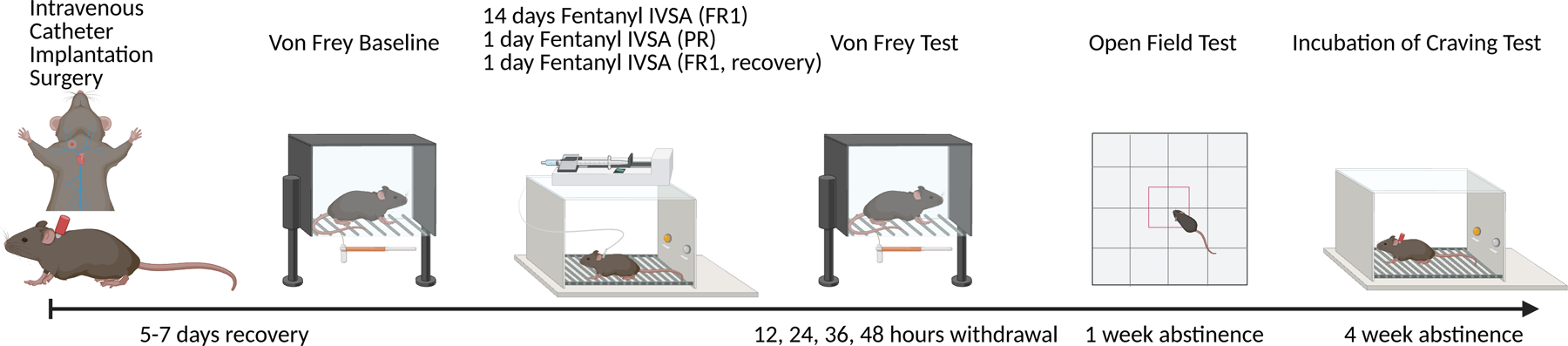
Experimental Design. Mice first underwent jugular catheterization surgery, followed by 5-7 days of recovery. Baseline mechanical pain sensitivity was then assessed using the von Frey test. Next, mice underwent 14 days of intravenous fentanyl self-administration (IVSA) followed by Progressive ratio (PR) testing. After PR, an additional day of IVSA was performed. Mechanical pain sensitivity was then measured at 12, 24, 36, and 48 hours into withdrawal. An open field test was conducted 1 week into abstinence, and an incubation of craving test was performed 4 weeks into abstinence.

### Jugular catheterization surgery

For implantation of jugular catheters, mice were anesthetized, using isoflurane (3%-4%), and maintained under anesthesia (1.5%-2%) during surgery. After shaving and applying iodine and 70% ethanol solution, incisions were made in the mid-scapular and anteromedial to the forearm regions. A catheter approximately 6 cm in length (i.d.: 0.31 mm, o.d.: 0.64 mm; VWR, PA), was affixed to a vascular access button (Instech Laboratories, Inc., Plymouth Meeting, PA), was passed subcutaneously from the back to the neck incision. After isolating the right jugular vein, the vein was punctured with a 23-gauge needle; the catheter was inserted to the point of a silicone bead positioned 1.2 cm from the catheter tip. Blood was drawn to confirm the correct placement in the vein, and the catheter was secured in place with silk-thread sutures immediately above and below the silicone bead. To avoid clotting, the catheter was flushed with 0.02ml sterile heparinized (20 USP units/ml) saline. The catheter was capped with a protective aluminum cap (Instech Laboratories). Antibiotic ointment was applied to the catheter exit wounds on the animal’s back and forearm. All mice were allowed to recover for 5–7 days before behavioral testing. The catheters were flushed with 0.02–0.03 ml of heparinized saline (20 USP units/ml) before and after each self-administration session. For all mice, the patency of the catheters was confirmed during training and after the study using 15 mg/ml ketamine in saline. Rapid loss of muscle tone was interpreted as an indication of patency.

### von Frey Test

Following recovery from surgery, mice were tested for baseline mechanical pain sensitivity using the von Frey Test. Mice were later tested for mechanical pain sensitivity at 12, 24, 36, and 48 hours into withdrawal from fentanyl. For von Frey testing, mice were held in Plexiglas boxes on a custom-made elevated metal wire surface and habituated to the behavioral apparatus for 30 min to ensure stable readouts of mechanical pain sensitivity. Nylon monofilaments (Stoelting Co., Wood Dale, IL) of forces ranging from 0.008 to 2 g were applied to the hind paw using the simplified up-down method (SUDO) (39, 40). Starting with a mid-range force (0.16 g), the filament was applied to the mid-plantar surface of the hind paw for ten trials, then repeated with ascending or descending forces depending on the number of paw withdrawals. A total of five stimuli were allowed for each trial. The data is presented as the average of left and right hind paw responses.

### Intravenous fentanyl self-administration (IVSA)

After recovery from surgery and baseline von Frey testing, the mice underwent a daily fentanyl IVSA paradigm for two weeks. Mice self-administered fentanyl during two-hour sessions every day over 14 consecutive days. The self-administration chambers (Med Associates, St Albans, VT) were located inside a larger sound attenuating box (Med Associates). The self-administration chambers were equipped with two retractable levers; one lever was designated as active and the other as inactive throughout all sessions. At the beginning of each session, the levers were extended, and the house light within the chamber was turned on, indicating the availability of the reward. A press on the active lever by the mouse triggered the infusion pump to deliver a 15 µl infusion of fentanyl (2.5 µg/kg of fentanyl per infusion) from a 10 ml syringe. Concurrently, a cue light positioned above the active lever would light up, and the house light would turn off. Following each infusion, there was a 20-second ‘time-out’ phase, during which responses to lever presses were logged but did not trigger programmed effects (e.g., drug infusion, cue light), and the house light would be off. Once the time-out was over, the house light would turn back on to indicate availability of reward again. Interactions with the inactive lever were also recorded, with no programmed consequences. For the mice that failed to press the active lever (0 active lever presses) in the first 2 hours of the first IVSA session, the session was extended until one active lever press was made or for a maximum of six hours. This extended session was given to ensure sufficient learning of the association between pressing the active lever and receiving drug reward occurred on the first day.

### Open Field Test

The Open Field Test was conducted when mice were 1-week into abstinence from fentanyl. All animals were acclimatized to the testing room for at least 30 minutes. Each mouse was then gently placed in the center of the open field apparatus (100 diameter) and allowed to explore freely for a 15-minute session. The sessions were recorded using an overhead camera connected to a computer running behavioral tracking software (Ethovision). The distance traveled was recorded to examine locomotion. The time spent in the predefined center area and the number of entries into the center zone were used to measure anxiety-like behavior.

### Incubation of Craving Test

Four weeks after the final IVSA session, an incubation of craving test was performed. The same procedural conditions as the previous IVSA sessions were used but without active lever presses providing drug reinforcement. Presses on the active lever activated the cue light, mirroring the conditions of drug availability, but no drug infusion was delivered. This procedure aimed to assess the persistence of drug-seeking behavior into a period of extended abstinence.

### Data Analysis

The results are expressed as mean ± SEM. The data were analyzed using analysis of variance (ANOVA), mixed effect analysis, or t-test as appropriate. The ANOVAs were followed by the Bonferroni *post hoc* test as appropriate. Data with missing values were analyzed by fitting a mixed model rather than repeated measure ANOVA, which cannot handle missing values.

## Results

### Male and female mice significantly escalate fentanyl self-administration over 14 days of access

Following recovery from surgery, male and female mice underwent 14 days of 2-hour fentanyl IVSA sessions. Inactive versus active lever presses were first compared to determine if mice showed significant discrimination of the active lever. Due to a programming error, inactive lever presses of 7 mice on day 1 were not recorded. These data were analyzed by fitting a mixed model rather than repeated measures ANOVA, which cannot handle missing values. Mixed effects analysis revealed a significant main effect of days (F_13, 221_ = 5.13, *p* < .001, lever type (F_1, 17_ = 55.65, *p* < .001), and their interaction (F_13, 213_ = 7.45, *p* < .001). The Bonferroni *post ho*c test showed that mice significantly increased fentanyl active lever presses starting on day 4 through day 14 compared to day 1 (active lever presses on day 1: 4.83 ± 1.38, and day 4: 30.00 ± 5.43). The active lever presses were significantly higher than inactive lever presses starting on day 5 through day 14 (Fig. 2A; Day 5 active lever presses: 34.67 ± 5.21, and inactive lever presses: 8.5 ± 1.43).

**Figure 2.**
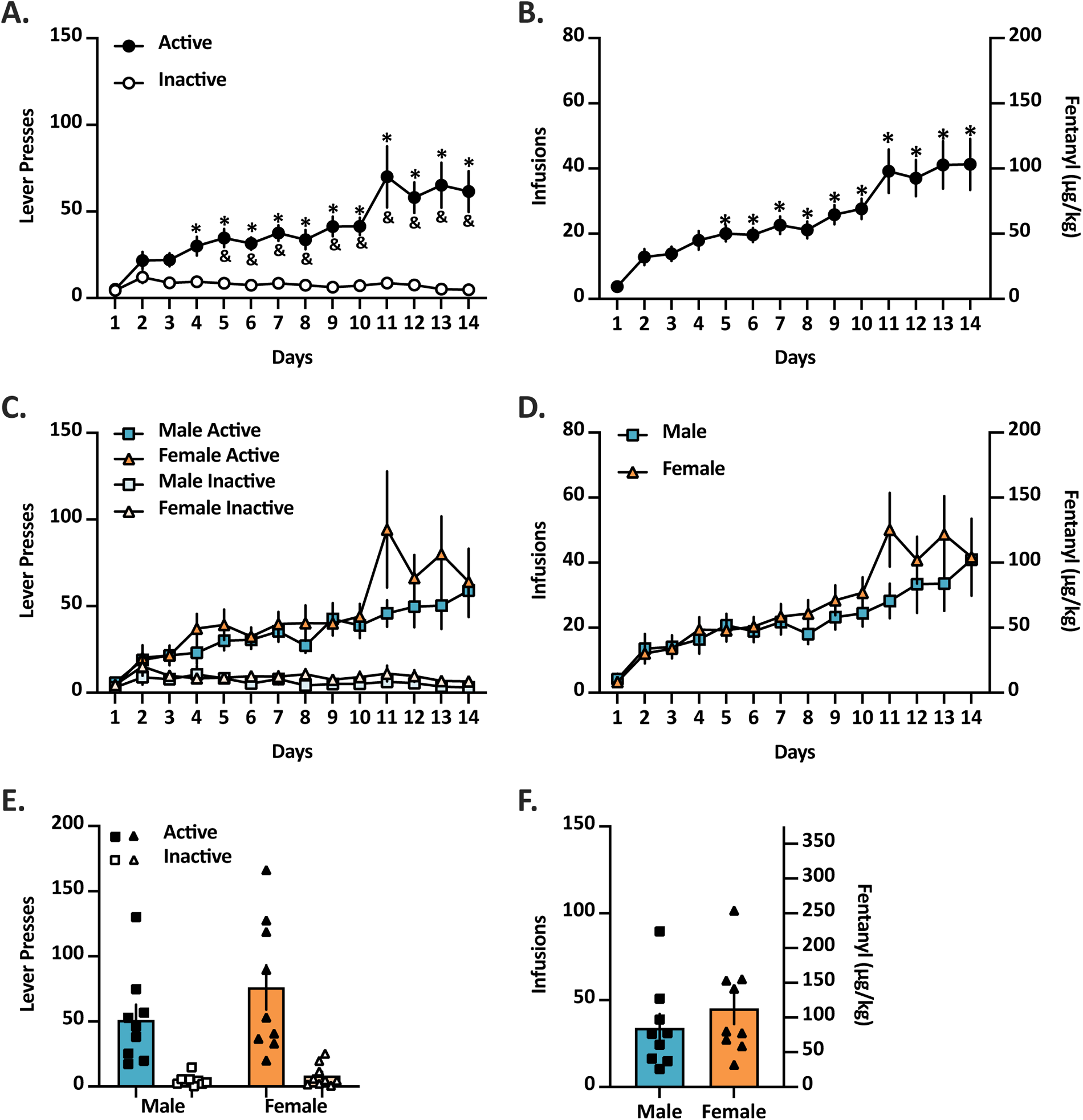
Fentanyl infusions and lever presses over 14 days of IVSA. **(A)** Mice pressed the active lever (black) significantly more than the inactive lever (white) starting on day 5 and continuing throughout the remaining sessions until day 14(*). Mice pressed the active lever significantly more than day 1 starting on day 4 and throughout day 14 (^&^). **(B)** Daily fentanyl infusions received (2.5 µg/kg/infusion) over 14 days of IVSA for all mice. Mice intravenously self-administered significantly more fentanyl starting on day 5 and continuing throughout the remaining sessions until day 14. **(C)** Active (dark blue and dark orange) and inactive (light blue and light orange) lever presses of male (blue square) and female (orange triangle) mice over 14 days of fentanyl IVSA. No sex difference was found between male and female lever pressing. **(D)** Daily fentanyl Infusions received over 14 days of IVSA of male (blue square) and female (orange triangle) mice. No sex difference was found between male and female infusions received. **(E)** Average active (black square and black triangle) and inactive (white square and white triangle) lever presses over the last 4 days of fentanyl IVSA (Day 11 to Day 14) in male (blue) and female (orange) mice. No sex difference was found between males and females over the last 4-day average lever presses. **(F)** Average fentanyl infusions received over the last 4 days of sessions in male (blue) and female (orange) mice. No sex difference was found between males and females over the average of the last 4-days of infusions received. Data are shown as mean ± SEM. n = 18; 9 males, 9 females. \**p* < .05 compared to day 1. ^&^*p* < .05 compared to inactive lever.

When the fentanyl infusions received were analyzed, repeated measures one-way ANOVA revealed a significant main effect of days of fentanyl infusions (F_13, 221_ = 10.86, *p* < .001). The Bonferroni *post hoc* test indicated that mice had significantly higher fentanyl infusions from day 5 through day 14 compared to day 1 (Fig. 2B; 3.78 ± 1.00 infusions on day 1 *vs.* 20 ± 2.40 infusion on day 5).

These data were further analyzed to determine if sex differences existed. When analyzing active versus inactive lever presses, with a main effect of sex, and lever type as the between-subject factor, there was no significant effect of sex found. There was a significant main effect of days (F_13, 208_ = 5.20, *p* < .001), lever type (F_1, 16_ = 55.29, *p* < .001), and interaction between days and lever type (F_13, 201_ = 7.68, *p* < .001), but no effect of sex, interaction between days and sex, interaction between lever and sex, or interaction between days, lever, and sex (Fig. 2C). When analyzing infusions received there was also no significant effect of sex found. The two-way repeated-measures ANOVA of sex and days revealed a significant main effect of days (F_13, 208_ = 10.82, *p* < .001) but not sex, or interaction of days and sex, on fentanyl infusion (Fig. 2D). The average of the last 4-days of lever presses and infusions of male and female mice were then compared. For the average of the last 4 days of lever presses, two-way ANOVA of sex and lever type revealed a significant effect of lever type (F_1, 16_ = 31.65, *p* < .001) but no effect of sex or interaction (Fig. 2E). For the average of the last 4 days of infusions, unpaired t-test revealed no significance between males and females (Fig. 2F).

### Male and female mice take significantly more fentanyl in the first 30 minutes of the 2-hour self-administration sessions after escalation of intake

To further analyze the self-administration data over the course of the 2-hour period once escalation of intake had been established, the average infusions received per 30-minute quarter from the last 4 sessions of fentanyl self-administration was examined. Repeated measures one-way ANOVA revealed a significant main effect of time quarter on fentanyl infusions (F_3, 51_ = 6.56, *p* < .001). The Bonferroni *post hoc* test indicated that mice received significantly more fentanyl infusions in the first 30 minutes (12.28 + 1.64) of the two-hour session compared to the second (9.05 ± 1.34), the third (9.40 ± 1.78), or the fourth (8.92 ± 1.68) 30 minutes blocks (Fig. 3A). The data were further analyzed, and no significant effect of sex was found. Repeated two-way ANOVA revealed significant effect of time (F_3, 48_ = 6.37, *p* = .001), but no effect of sex, or interaction (Fig. 3B).

**Figure 3.**
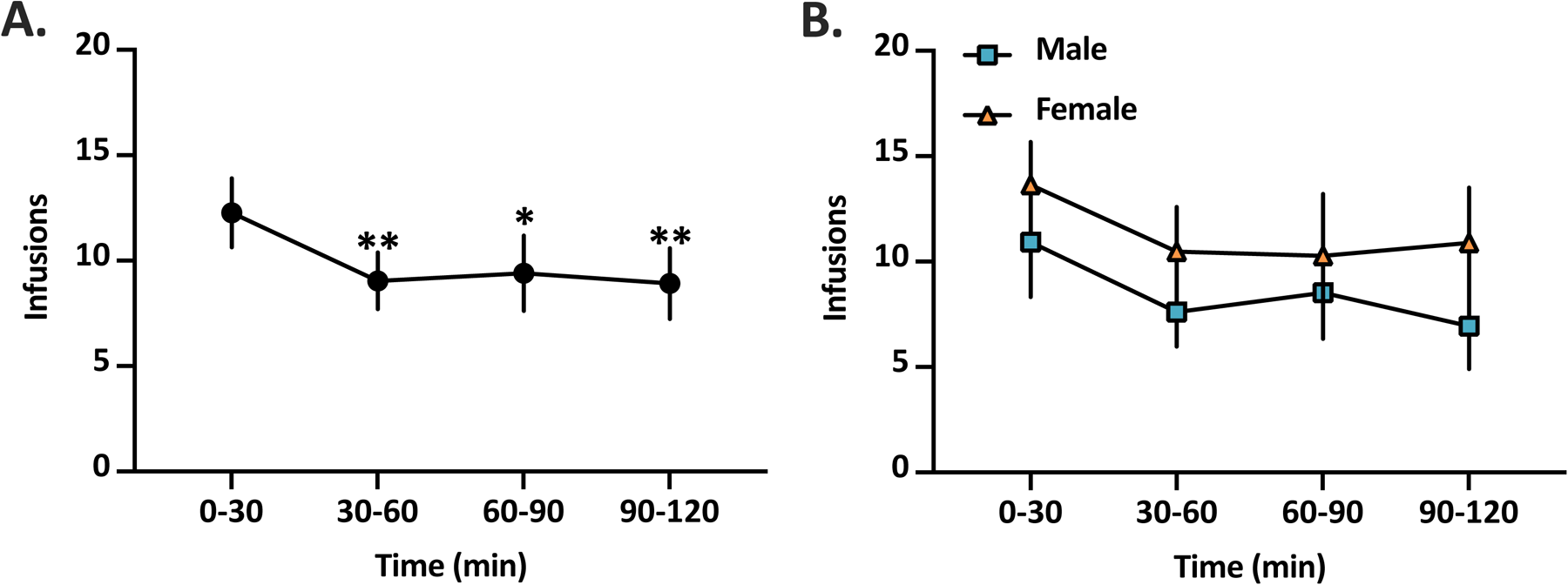
Average infusions received per 30-minute quarter from the last 4 sessions of fentanyl IVSA. **(A)** Average fentanyl infusions received from the last 4 sessions of fentanyl IVSA of all mice. Mice received significantly more fentanyl infusions during *the* first *30* minutes of the sessions compared to the subsequent three 30-minute intervals. **(B)** Average fentanyl infusions received from the last 4 sessions of fentanyl IVSA in male (blue square) and female (orange triangle) mice. No significant differences in infusions were observed between male and female mice across the four 30-minute intervals. Data are shown as mean ± SEM. n = 18; 9 males, 9 females. \**p* < .05; \*\**p* < .01 compared to infusions in 0-30mins interval

### Male and female mice show increased mechanical pain sensitivity during fentanyl withdrawal

The von Frey test was performed at multiple timepoints during withdrawal to determine if fentanyl withdrawal would result in increased sensitivity to mechanical pain. Repeated measures one-way ANOVA revealed a significant main effect of time of withdrawal on paw withdrawal threshold (F_4, 48_ = 13.82, *p* < .001). The Bonferroni *post hoc* test indicated that mice showed a significantly decreased paw withdrawal threshold, indicating increased mechanical pain sensitivity, at 36-, and 48-hour withdrawal when compared to baseline measurements (Fig. 4A). The data were then analyzed with sex as the between subject factor and no effect of sex was found. Repeated two-way ANOVA revealed significant effect of time of abstinence (F_4, 44_ = 14.21, *p* < .001), but no effect of sex, or interaction (Fig. 4B).

**Figure 4.**
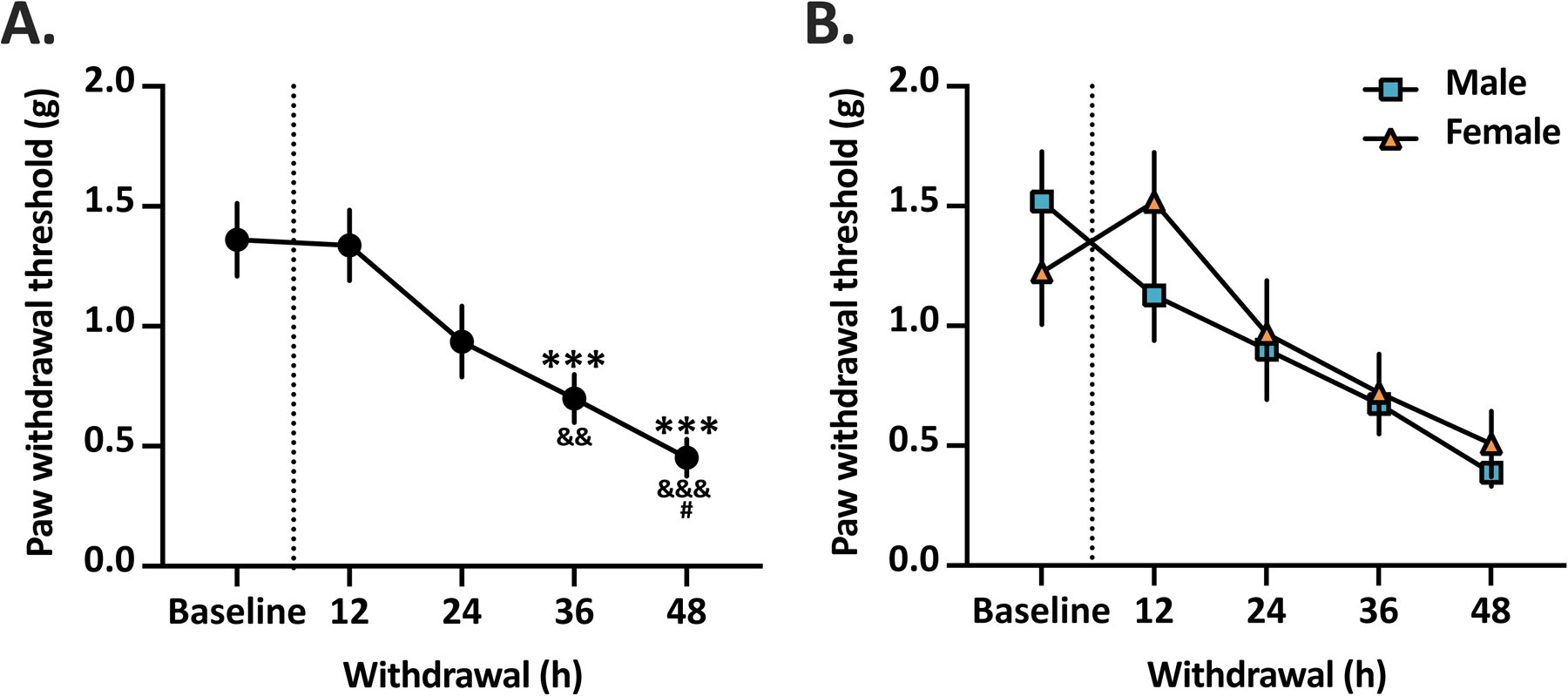
von Frey test during fentanyl withdrawal. **(A)** Paw withdrawal threshold of mice at baseline and 12-, 24-, 36-, and 48-hourfentanyl withdrawal. Mice showed a significant decrease in paw withdrawal thresholds at 36-, and 48-hour withdrawal compared to their own baseline measurement (*). Mice showed a significant decrease in paw withdrawal threshold at 36- and 48-hour withdrawal compared to 12-hour withdrawal (^&^). Mice showed a significant decrease in paw withdrawal threshold at 48-hour withdrawal compared to 24-hour withdrawal (^#^). **(B)** Mechanical pain sensitivity of male (blue square) and female (orange triangle) at baseline and 12, 24, 36, and 48 hours into abstinence. No significant difference was found between male and female mice for paw withdrawal threshold during fentanyl withdrawal. n = 13; 6 males, 7 females. Data are shown as mean ± SEM. \*\*\**p* < .001 compared to baseline ^&&^*p* < .01, ^&&&^*p* < .001 compared to 12-hour withdrawal. ^#^*p* < .01, compared to 24-hour withdrawal.

### Male and female mice show anxiety-like behavior 1 week into abstinence from fentanyl

To determine if 1 week of abstinence from fentanyl would produce anxiety-like behavior, the open field test was performed. When analyzing center time, unpaired t-test revealed a significant difference in time spent in the center area (t_26_ = 2.73, *p* = .011) between mice that were 1 week into abstinence from fentanyl and naive mice. Mice 1 week into abstinence from fentanyl spent significantly less time in the center area of the open field (101.1 ± 8.75 s) compared to naive mice (162.6 ± 18.92 s; Fig. 5A). When analyzing number of entries to the center area, unpaired t-test revealed a significant difference in number of entries to the center area (t_26_ = 2.90, *p* = .008). Mice 1 week into abstinence from fentanyl entered the center area with fewer numbers (59.92 ± 4.55 entries) than naive mice (81.53 ± 5.72 entries; Fig. 5B). There was no significant difference in in distance traveled between groups (Fig. 5C).

**Figure 5.**
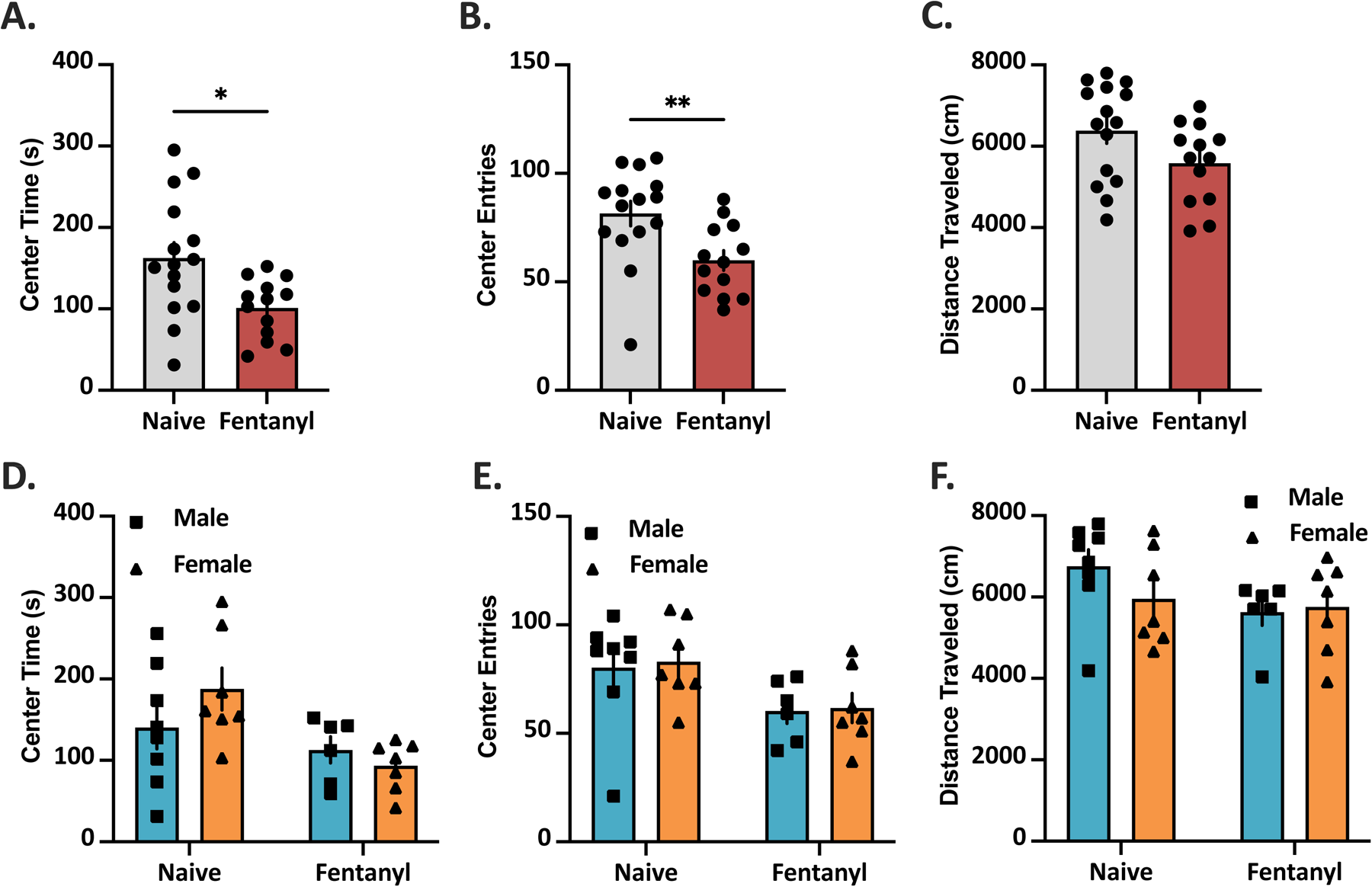
Open field test during abstinence from fentanyl. **(A)** Time spent in the center area (s), **(B)** frequency to enter the center area, and **(C)** distance traveled (cm) in the open field test of naive mice and mice 1 week into abstinence from fentanyl. During abstinence from fentanyl, mice spent significantly less time in the center area and made fewer entries into the center area than naive mice. Abstinence from fentanyl did not affect locomotor behavior. **(D)** Time spent in the center area (s), **(E)** frequency to enter the center area, and **(F)** distance traveled (cm) in the open field test of male (blue) and female (orange), naive mice, and mice 1 week into abstinence from fentanyl. No significant sex difference was found. Naive: n = 15; 8 males, 7 females; Fentanyl abstinence: n = 13; 6 males, 7 females. *p < .05, **p < .01

When separately analyzing the data for sex differences, no effect of sex was found. For center time, two-way ANOVA revealed an effect of abstinence from fentanyl (F_1, 24_ = 7.68, *p* = .011), but no effect of sex or interaction (Fig. 5D). For entries to the center area, two-way ANOVA also revealed a significant effect of abstinence from fentanyl (F_1, 24_ = 7.26, *p* = .013), but no effect of sex or interaction (Fig. 5E). Neither sex nor abstinence from fentanyl had an effect on distance traveled of the mice (Fig. 5F).

### Male and female mice show drug-seeking behavior 4 weeks into abstinence from fentanyl

To assess whether a craving for fentanyl would persist four weeks into abstinence, mice were subjected to a two-hour IVSA procedure identical to their previous sessions, except no drug was infused upon pressing the active lever. A comparative analysis between the number of active lever presses on the 14th day of self-administration and the day of the craving test was conducted. Using a paired t-test, a significant increase in active lever presses was observed during the craving test as compared to the last day of IVSA (t_9_ = 3.75, *p* = .005).

Further analysis was carried out to explore potential sex differences in craving behaviors. A two-way repeated measures ANOVA was applied, which identified a significant main effect of the four-week abstinence on active lever presses (F_1, 8_ = 12.50, *p* = .008). However, there was no significant effect based on sex, nor was there a significant interaction between sex and the abstinence period (Fig. 6B).

**Figure 6.**
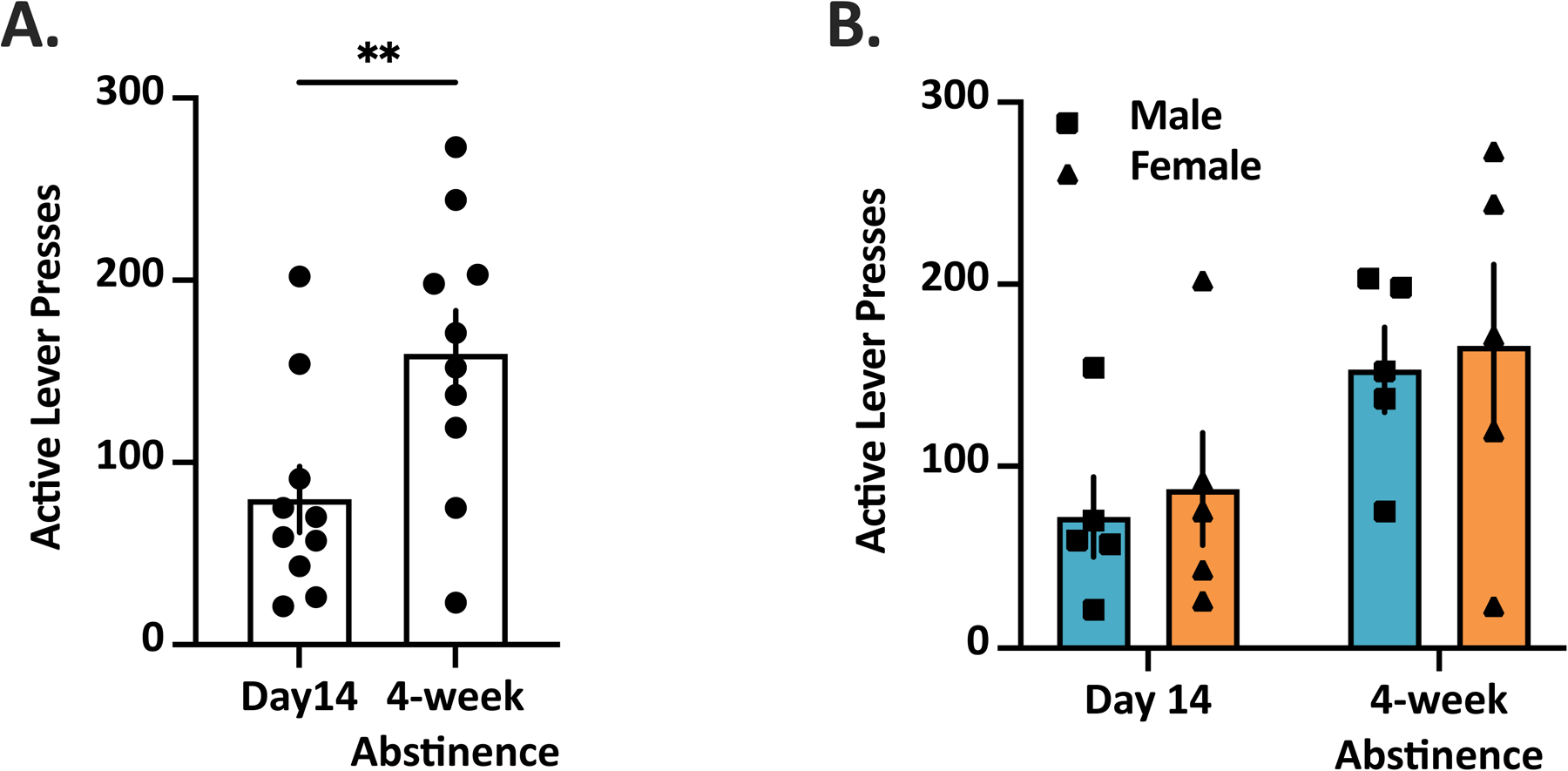
Incubation of craving test. **(A)** Active lever presses of mice at day 14 of IVSA and 4 weeks into abstinence. Mice showed a significant increase in active lever presses at craving test day compared to day 14 of IVSA. **(B)** Active lever presses of male (blue square) and female (orange triangle) at day 14 of IVSA and 4 weeks into abstinence. No significant sex difference was found for active lever presses of mice on day 14 IVSA and 4 weeks into abstinence. n = 10; 5 males, 5 females. Data are shown as mean ± SEM. \*\**p* < .01.

## Discussion

The current study aimed to develop a translationally relevant mouse model of fentanyl use disorder (FUD) by employing chronic fentanyl intravenous self-administration (IVSA). Throughout the experiment, male and female mice displayed no significant differences and both groups exhibited a substantial escalation in their fentanyl intake over the 14-day period. Mice demonstrated front-loading behavior, consuming fentanyl more rapidly during the first 30 minutes of each self-administration session. Notably, mice exhibited a significant increase in pain sensitivity during the 36-to 48-hour period following fentanyl withdrawal and displayed anxiety-like behavior one week into fentanyl abstinence. Mice additionally showed persistent craving for fentanyl four weeks into abstinence. Collectively, these results establish a mouse model of FUD that encompasses key characteristics of fentanyl dependence, including the escalation of fentanyl intake, front-loading behavior, the manifestation of significant withdrawal symptoms, and incubation of craving.

The present study found that mice significantly escalated their fentanyl intake over the course of two weeks of IVSA of fentanyl. Interestingly, in this study, mice were found to take more fentanyl during the first 30 minutes of the two-hour session than other portions of the session, implying front-loading behavior. Front-loading behavior, which refers to the rapid consumption of a drug at the beginning of a self-administration session, has been more investigated in alcohol drinking (41–43). In the present study, the “loading” phase observed within the first 30 minutes may indicate an increased drug intake due to the emergence of negative emotional-like states associated with withdrawal. This behavior aligns with the theory that negative reinforcement mechanisms drive excessive drug intake to alleviate discomfort from withdrawal (3, 44, 45) and is consistent with previous findings in a rat model of oxycodone self-administration that observed front-loading behavior (46).

The current study builds upon previous findings and demonstrates that 2 hours of access to 2.5 µg/kg/infusion of fentanyl over 14 days is sufficient to establish escalation of fentanyl intake and result in withdrawal symptoms in mice. Previous literature has investigated rodent models of opioid dependence, including morphine (47–49), oxycodone (10, 12, 50, 51), heroin (6, 52–54), remifentanil (34, 55–58), and fentanyl (24, 33, 36, 38, 59–66), using various routes of administration, with the majority of studies done in rats. IVSA of fentanyl has been widely studied in rats (24, 25, 59–62, 67–71), with findings demonstrating mixed results on the development of dependence and withdrawal signs. One previous study reported that rats undergoing 14 days IVSA of fentanyl for 6 hours per day maintained a consistent rate of self-administration (62). Another study found that rats self-administering fentanyl for 3 hours per day over 10 days exhibited a trend towards increasing reward-associated lever presses (59). A separate study showed that when compared to a short access self-administration period, 12-hour self-administration of fentanyl resulted in escalation of fentanyl intake (25). In addition to rats, mouse models of fentanyl self-administration have been developed (33, 63, 66, 72, 73), although fentanyl IVSA in mice remains relatively understudied. One prior study demonstrated that mice maintained stable levels of oral fentanyl consumption and showed a significant preference for the active lever associated with fentanyl in an operant chamber (63). Another study utilized a drinking in the dark paradigm to model oral fentanyl self-administration in mice, finding that mice consumed increasing amounts of fentanyl as the dose increased (37). Leonardo et al. (33) successfully established fentanyl IVSA in mice, showing rapid acquisition and a dose-responsive curve.

In the present study, a significant increase of mechanical pain sensitivity starting 36 hours after the cessation of fentanyl use, persisting up to 48 hours (and potentially longer) into withdrawal was observed. Additionally, increased anxiety-like behavior was found one week into abstinence from fentanyl. Withdrawal from opioids such as heroin and morphine has been shown to result in hyperalgesia in humans (74, 75) and similarly in rats and mice (16, 18). Recent studies have shown oxycodone withdrawal to result in increase of mechanical pain sensitivity in rats (10) and somatic withdrawal signs in mice (20–22). Overall, these prior studies are consistent with the current results from withdrawal from IVSA of fentanyl, suggesting similar withdrawal symptoms across opioids in preclinical studies. Fentanyl withdrawal has been investigated using osmotic pump implantation in rats, with researchers finding significant increases in somatic withdrawal signs of fentanyl withdrawal rats compared to control rats (76–78). Consistent with the increased anxiety-like behavior observed 7 days into abstinence in the current study, a previous study found decreased time in the open arms of the elevated plus maze in mice after 10 days of fentanyl abstinence (38). Another study found female mice spent decreased time in the center of the open field 6 weeks post-morphine withdrawal (47), suggesting the effects observed in the current study may persist even longer. Affective withdrawal symptoms have been suggested to be the main contributors to substance use disorder rather than somatic withdrawal signs (79, 80). Finding of increased mechanical pain sensitivity and increased anxiety-like behavior in the current study suggests a strong relevance of the IVSA of fentanyl model to studying FUD. Together with findings from previous literature, these results contribute to the understanding of opioid withdrawal, especially fentanyl withdrawal, which may have substantial implications for treatment approaches in humans (81).

The present study investigated the impact of a four-week abstinence period on drug-seeking behavior in mice that had previously escalated intake of fentanyl. The results demonstrate that both male and female mice exhibited drug-seeking behavior following the abstinence period, as evidenced by a significant increase in active lever presses during an incubation of craving test compared to the last day of IVSA. These findings are consistent with previous studies that have reported the incubation of drug craving in rodents following a period of abstinence from various drugs of abuse (82–84). The current study extends these findings to fentanyl, providing an important measure for translational relevance. The observed drug-seeking behavior following a four-week abstinence period highlights the persistent nature of fentanyl-related craving and the potential risk of relapse even after an extended period of abstinence. The present study did not find significant sex differences in drug-seeking behavior following the abstinence period. The mechanisms underlying the incubation of fentanyl craving remain to be elucidated. Previous studies have implicated various neurobiological processes, such as changes in glutamatergic and dopaminergic neurotransmission, alterations in brain-derived neurotrophic factor signaling, and epigenetic modifications, in the development and expression of incubated drug craving (83, 85). Future studies can use the IVSA of fentanyl model to further explore the specific molecular and neural substrates involved in the incubation of fentanyl craving to identify potential therapeutic targets for relapse prevention.

This study provides a translationally relevant mouse model of fentanyl dependence, aligning closely with the criteria for translational value and utility in basic science (86). By facilitating voluntary drug self-administration, the model effectively mirrors the voluntary nature of human substance use. The manifestation of behaviors such as escalated drug intake over time, increased sensitivity to pain during withdrawal, the exhibition of anxiety-like behaviors and incubation of craving during abstinence, lend credence to the face validity of our model. These behaviors not only parallel the psychological and physiological complexities of human FUD but also underscore the model’s applicability in exploring the nuances of substance use disorder (44, 87). While our findings lay a robust foundation for understanding fentanyl dependence, they also highlight the potential for future research to enhance predictive validity. This could involve testing the efficacy of interventions known to mitigate human FUD, thereby bridging the gap between animal models and clinical applications. The current study provides a model that not only reflects the intricacies of human fentanyl dependence but also paves the way for future exploration into interactions with other drugs, neurobiological mechanisms underlying FUD and therapeutic treatments. For example, xylazine is commonly used in combination with fentanyl (88) and the two interact together (89). Xylazine has also recently been shown to have independent effects on kappa opioid receptors that may act to potentiate the effect of fentanyl (90).

In conclusion, the present study establishes a translationally relevant mouse model of FUD that recapitulates key features of the human condition, including escalation of drug intake, front-loading behavior, the presence of withdrawal signs, and incubation of craving. These findings provide valuable insights into the development and maintenance of fentanyl dependence and offer a powerful tool for further investigating the neurobiological underpinnings of this disorder. The present study addresses the current lack of mouse models that examine fentanyl withdrawal symptoms, particularly affective symptoms such as anxiety-like behavior. By incorporating these measures, a more complete representation of the FUD phenotype is provided. However, the model does not currently investigate other aspects of FUD, such as compulsive drug-seeking despite negative consequences or the impact of social and environmental factors on fentanyl use. Future studies could adapt our model to include these features, such as using punishment paradigms or social housing conditions, to further enhance its translational validity.

## Acknowledgements

This work was supported by the Purdue Women’s Global Health Institute and Ralph W. and Grace M. Showalter Research Trust.

