## Supplemental Information for "Escalation of intravenous fentanyl self-administration and assessment of withdrawal behavior in male and female mice"

**Supplemental Methods**

*Progressive Ratio (PR)*

In the PR test, the number of active lever presses required to obtain each successive infusion increased according to the following series: 1, 2, 4, 6, 9, 12, 15, 20, 25, 32, 40, 50, 62, 77, 95, 118, 145, 178, 219, 268, 328, 402, 492, 603, 737, 901, 1102, 1347, 1646, 2012 (88). The breakpoint was defined as the last completed ratio before a 1-h period during which no infusions were obtained. The PR test lasted for 2 hours.

Supplemental Results

*Male and female mice display comparable fentanyl-seeking behavior during PR test.*

Male and female mice exhibited similar levels of fentanyl-seeking behavior and motivation during the PR test following 14 days of fentanyl IVSA. No significant differences were observed between sexes in the number of active lever presses (Figure S1A) or the breakpoint (Figure S1B) achieved during the PR test.

*Mice maintain consistent fentanyl intake on recovery day prior to von Frey withdrawal*

Following the PR test, mice were given a single recovery day of IVSA to ensure that drug intake was similar to levels comparable to those observed on the last day of the 14-day IVSA period (Figure S2A) before the withdrawal tests. Both male and female mice exhibited similar levels of fentanyl self-administration on the recovery day compared to Day 14 of the IVSA period (Figure S2B).

Figure S1. F
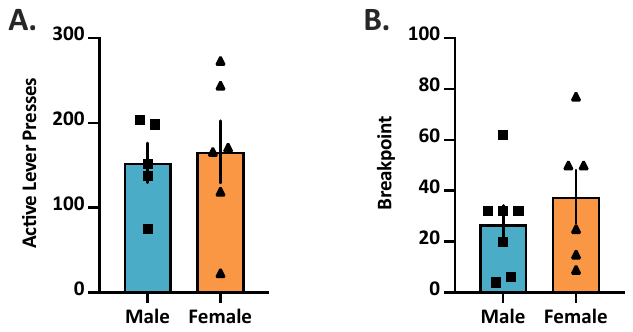
entanyl-seeking behavior and motivation during progressive ratio test following 14-day fentanyl IVSA in male and female mice. (A) Number of active lever presses by male (blue) and female (orange) mice during the PR test. (B) Breakpoint achieved by male and female mice during the PR test. Data are presented as mean ± SEM. No significant differences were observed between male and female mice. n = 11; 5 males, 6 females.

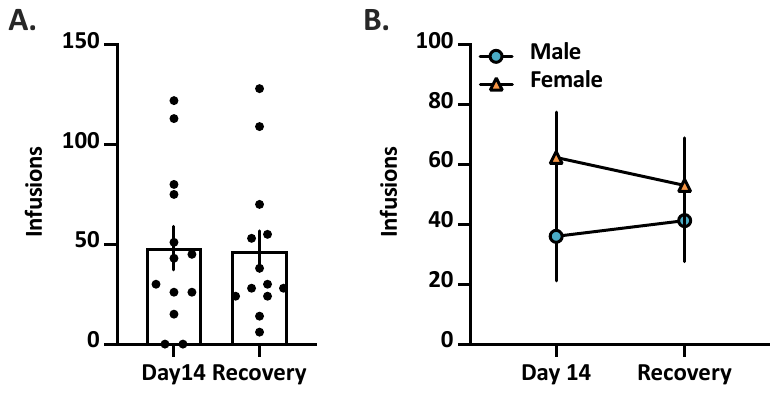

Figure S2. Fentanyl self-administration on Day 14 and the recovery day in male and female mice. (A) Number of fentanyl infusions earned by mice on Day 14 IVSA and on the recovery day. (B) Number of fentanyl infusions earned by male mice on Day 14 IVSA and on the recovery day. Data are presented as mean ± SEM. n = 13; 6 males, 7 females.
